# OncoRep: An n-of-1 reporting tool to support genome-guided treatment for breast cancer patients using RNA-sequencing

**DOI:** 10.1101/008748

**Authors:** Tobias Meissner, Fisch Kathleen M., Louis Gioia, Andrew I. Su

**Author notes:** These authors should be considered to have equal author status (Tobias Meissner, Fisch Kathleen M.).

## Abstract

Breast cancer comprises multiple tumor entities associated with different biological features and clinical behaviors, making individualized medicine a powerful tool to bring the right drug to the right patient. Next generation sequencing of RNA (RNA-Seq) is a suitable method to detect targets for individualized treatment. Challenges that arise are i) preprocessing and analyzing RNA-Seq data in the n-of-1 setting, ii) extracting clinically relevant and actionable targets from complex data, iii) integrating drug databases, and iv) reporting results to clinicians in a timely and understandable manner. To address these challenges, we present OncoRep, an RNA-Seq based n-of-1 reporting tool for breast cancer patients. It reports molecular classification, altered genes and pathways, gene fusions, clinically actionable mutations and drug recommendations. It visualizes the data in an approachable html-based interactive report and a PDF clinical report, providing the clinician and tumor board with a tool to guide the treatment decision making process. OncoRep is free and open-source, thereby offering a platform for future development and innovation by the community.

## Introduction

Breast cancer is the leading cause of cancer among females making up 23% of total cancer deaths^1^. It is a heterogenous disease comprising multiple tumor entities associated with distinctive histological patterns, different biological features and clinical behaviors^2, 3^. This is driven by the fact that different breast cancer subtypes are characterized by distinct molecular, genetic, epigenetic, and transcriptional patterns (e.g. gene amplifications, in-frame fusion genes or mutations, homozygous deletions, disrupting fusions and deleterious mutations)^4^. Five year survival rates from the time of diagnosis range from 98 percent (localized cancer) to 24 percent (metastatic cancer). Twenty percent of patients who completed either adjuvant or neoadjuvant systemic therapy had a recurrance of the disease within 10 years after treatment ^5, 6^

Molecularly profiling breast cancer tumors takes advantage of the genomic characteristics of the tumor to improve the chances of patient response to targeted agents. This enables stratification of patients based on their molecular alterations. Therapies targeting specific genomic alterations have been shown to be effective in treating specific subgroups of breast cancer patients. Examples of targeted therapies include the efficacy of Trastuzumab in *HER2*-amplified breast cancers, the mTOR inhibitor Everolimus in hormone receptor positive, *HER2*-negative patients, and the PARP inhibitor Olaparib in patients whose tumors harbor *BRCA1/2* mutations^7–10^ However, the transition to an individualized medicine approach, in which one selects the optimal treatment for a patient based on genomic information remains challenging. One of the main challenges is the translation of tumor genome-based information into clinically actionable findings. This relies not only on the identification of biologically relevant alterations that can be used as therapeutic targets or predictive biomarkers^4^, but also on the availability of appropriate reporting tools. These reporting tools need to integrate the wealth of genomic data and make it usable in a routine clinical setting. This will provide additional treatment options based on the genetic nature of the patient’s tumor, enabling true individualized cancer medicine.

Gene expression profiling using RNA-sequencing (RNA-Seq) is an ideal tool to assess the molecular heterogeneity of breast cancer to inform individualized medicine. It enables the estimation of transcript abundance, the detection of altered genes and molecular pathways, the detection of fusion genes and the reliable identification of genomic variants ^11–15^. RNA-Seq can be performed for nearly all breast cancer and metastatic breast cancer patients that require therapy using tissue collected during routine biopsy. The main difficulties remaining for prospective use of RNA-Seq in individualized breast cancer treatment are analyzing RNA-Seq data in the n-of-1 setting and the lack of an open source reporting tool providing clinically actionable information.

To address these challenges, we developed OncoRep, an open-source RNA-Seq based reporting framework for breast cancer individualized medicine https://bitbucket.org/sulab/oncorep. It can be used as part of the reproducible, automated next generation sequencing pipeline Omics Pipe^16^, it can be used as a standalone reporting tool and it can be adapted to existing sequencing pipelines. OncoRep includes molecular classification, detection of altered genes, detection of altered pathways, identification of gene fusion events, identification of clinically actionable mutations (in coding regions) and identification of target genes. Furthermore, OncoRep reports drugs based on identified actionable targets, which can be incorporated into the treatment decision making process. To demonstrate the feasibility of OncoRep, we produced reports based on the mRNA profiles of 17 breast tumor samples of three different subtypes (TNBC, non-TNBC and HER2-positive) which have been previously analysed and described^17–19^.

## Results

OncoRep was integrated as an RNA-seq Cancer Report pipeline in Omics Pipe^16^ which handles the processing of the raw RNA-seq data in an automated and parallel manner on a compute cluster. After the data were processed, the results files from each step and the patient specific meta data were automatically processed by OncoRep to produce a summary report for each patient. OncoRep performs the following analyses (Figure 1): i) variant annotation; ii) gene expression estimation; iii) differential gene expression analysis; iv) pathway analysis; v) prediction of receptor status and molecular subtype; and vi) selection of drugs targeting dysregulated genes, variants and pathways. OncoRep displays these results along with the results from the quality control of the raw data and alignment, variant calling, fusion gene detection and estimation of oncogenic potential. The R package knitr is used to produce an interactive HTML report. A PDF file containing a final summary report is generated using the R package Sweave (Figure 2). Analyzing a single patient sample (20-30 mio reads, 100bp, paired end) takes about one day in a cluster environment using four nodes.

**Figure 1.**
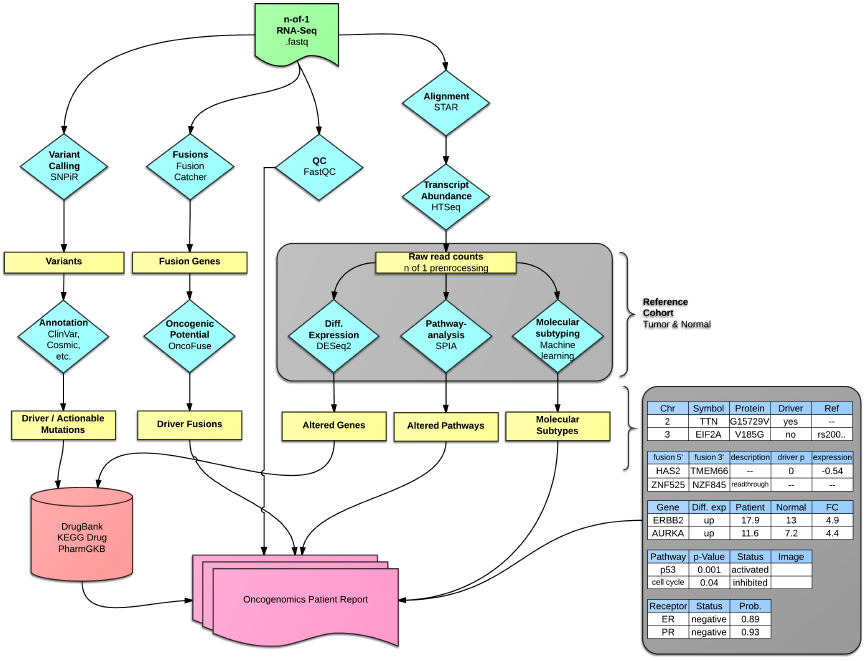
Flowchart illustrating tools used and their interactions within OncoRep. The four main branches (left to right) are variant calling, fusion gene detection, quality control and gene expression quantification and analysis (for a detailed description of each step see materials and methods). Results from each branch are analyzed, annotated and integrated and an HTML report is created at the final stage of the pipeline.

**Figure 2.**
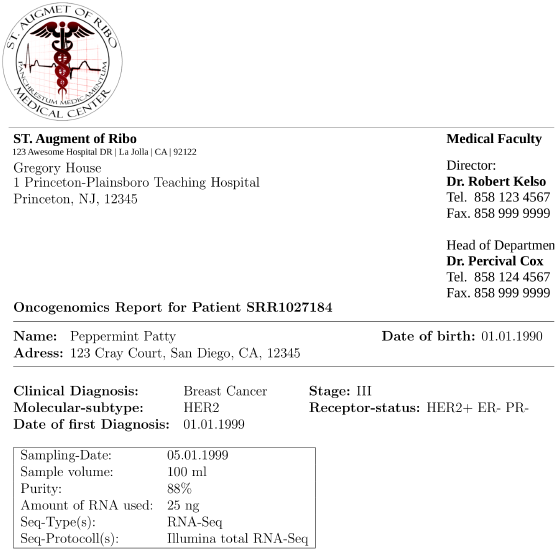
PDF clinical report generated by OncoRep for dissemination to treating physicians.

### Interactive Report

The HTML report produces interactive tables that are sortable and searchable. They can be exported as CSV files to be viewed in spreadsheet software. Gene descriptors and drugs are linked to the respective databases for easy access to further information. Pathways are visualized and they are annotated with differentially expressed genes. The interactive HTML reports for the 17 analyzed breast tumor samples can be viewed and browsed at http://sulab.org/tools/oncorep-oncogenomics-report/.

### PDF Report

The PDF based report is generated in `LATEX`, making it fully customizable (Figure 2). The report, as displayed here, holds basic patient information, sample processing information and gives a list of FDA approved drugs recommended based on the altered variants, genes and pathways in a patient’s tumor. An appendix holds all results from the various analysis steps in tabular form.

### Quality control

OncoRep provides quality control of raw RNA-Seq reads using the FastQC tool. Basic QC results are displayed within the HTML report and linked to the detailed FastQC report for further inspection if needed (for details see online Materials and Methods). Post alignment QC includes computation of insert size distribution and collecting basic RNA-Seq metrics using functionalities provided by Picard tools. The QC results and figures are presented within OncoRep.

### Variant Calling

Variants identified using the SNPiR pipeline^15^ are provided in a tabular format in the HTML report. If available, the user is displayed with clinically relevant information on the variants (e.g. a matching drug or the NCBI ClinVar rating). The variants are annotated using information from SnpEff^20^, dbNSFP^21^, COSMIC^22^, NCBI ClinVar^23^, CADD^24^, DrugBank^25^, PharmGkb^26^ and IntOGen^27^ (for details see online Materials and Methods). Furthermore, variants are matched against SNP-drug relationships available from DrugBank and PharmGkb and possible hits are displayed in the table.

### Fusion Gene Detection

Identified fusion gene candidates are provided in tabular manner in the HTML report. The information provided includes 5’ and 3’ fusion partners, fusion description (if available), and the the oncogenic potential prediction depicted as a p-value and expression gain/loss (for details see online Materials and Methods).

### Differential Gene Expression

OncoRep filters out all genes estimated to have ‘unreliable expression’ based on the expression of a background gene set of 156 genes that are not expressed in any sample of the reference cohort (see online Materials and Methods). All remaining genes are further analyzed. Differentially expressed genes are detected by comparing the reliably expressed genes in the patient tumor to normal breast tissue samples. The results are presented in tabular format in the HTML report.

### Pathway Analysis

Pathway analysis is conducted based on the differential expressed genes. Altered pathways are presented in tabular form in the HTML report. Visualizations of the pathways are provided with the differentially expressed genes colored based on their log2FoldChange expression compared to normal tissue.

### Receptor Status

OncoRep includes predictors for the three receptors ER, PR and HER2 (see online Materials and Methods for details). A new patient sample is classified as being positive or negative for the expression of each receptor and the prediction probability is given. Results are presented in tabular format in the HTML report.

### Molecular Subtype

OncoRep includes a predictor for the molecular subtype of the sample (Basal, HER2, Luminal A and Luminal B). A new patient sample is classified into one of the groups and the prediction probability is given. Results are presented in tabular manner in the HTML report.

### Drug Matching

OncoRep reports FDA approved compounds that target the discovered differentially expressed genes, variants and pathways in the patient sample. Results are presented in tabular manner in the HTML report. Results are linked to their DrugBank and KEGG Drug entries for further investigation.

## Discussion

In this article, we introduce OncoRep, a reporting tool that performs automated processing and interpretation of RNA-Seq raw data from breast cancer patients. Gene expression profiling using RNA-Seq generates vast amounts of data. This requires precise analyses and expert knowledge to generate clinically actionable information. Without expert knowledge, it remains challenging and time-consuming to do even simple data preprocessing and analysis. In a clinical setting, only clinically relevant data are needed from the RNA-Seq data. We address this problem by chaining software tools together to integrate them into a single analysis workflow that is able to deliver clinically digestable information within a short time span. OncoRep enables the prospective use of transcriptomic profiles within a clinical setting by performing molecular profiling, assessing altered genes and pathways, identifying mutations and fusion gene transcripts and by providing drug recommendations based on actionable targets to guide the treatment decision making process. This represents a critical first step towards individualized cancer treatment since it provides a reproducible approach in reporting actionable targets and allows for a quick turnaround time for real-time treatment of patients.

OncoRep detects altered genes, variants, fusions and dysregulated pathways in a patient’s tumor. The challenge exists to distill this large amount of information into clinically actionable targets. OncoRep draws from several databases and employs several variant filtering and annotation steps to extract variants that are the most biologically meaningful. Integrating these databases and presenting them in a report provides the community with a valuable resource, as many databases are sparsely populated and information is distributed throughout many poorly curated databases and in the primary literature^28^. OncoRep also reports fusion genes annotated with their predicted oncogenic potential, as many fusion genes have been discovered in breast cancer that may make a substantial contribution to its development ^14, 29, 30^. OncoRep uses several lines of molecular evidence to match drugs to altered drug targets in a patient’s tumor by drawing on information provided by DrugBank, KEGG Drug and PharmGKB.

By distilling and reporting clinically actionable aberrations on an individual level, OncoRep provides researchers and clinicians with a powerful tool for implementing individualized medicine.

For example, an OncoRep report for a patient may detect an aberration that is present in a small fraction of patients (e.g *ROS1* expression) for which targeted therapies exist. Since these are found in only a small fraction of patients, these treatments would not be used as standard of care, highlighting the importance of this method for identifying individualized treatments. In addition, OncoRep reports fusion genes and evidence exists that fusion genes may be suitable therapeutic targets. For example, Banerji *et al.* identified a recurrent *MAGI3-AKT3* fusion enriched in triple-negative breast cancer that leads to constitutive activation of AKT kinase, which can be targeted with an ATP-competitive AKT small-molecule inhibitor^29^. OncoRep advances individualized medicine by reporting all relevant information in a user-friendly way so that clinicians can access all of the results, as well as by extracting clinically actionable findings to aid in the treatment decision making process.

OncoRep overcomes one of the main difficulties remaining for prospective use of transcriptome profiling in clinical routine by creating reproducible and clinically digestible reports to guide clinical decision making. OncoRep is an open-source project, which increases the reproducibility and transparency of the analyses. We invite researchers to use the code, refine it and provide further improvements, such as incorporating new methods and additional disease areas. We believe that offering this modular and extensible framework will provide a useful community platform for implementing individualized genomic medicine.

## Methods

Methods and associated references are available in the online version of the paper.

## Online Methods

### Software design

OncoRep is developed within the open-source software environments R (v3.0.2)^31^ and Bioconductor (v2.13)^32^ using the knitr & knitr bootstrap packages for creating the patient report in HTML format and Sweave package for creating the PDF-based report. OncoRep is distibuted via Omics Pipe^16^ which handles the processing of the raw RNA-Seq data using distributed computing either on a local high performance cluster or on Amazon EC2. Installation and setup are documented online at http://pythonhosted.org/omics_pipe/.

### Reference cohort

The reference cohort incorporated into OncoRep (n=1,057) consists of 947 breast cancer samples and 106 matched tumor normal tissue samples from The Cancer Genome Atlas (TCGA), one normal breast tissue sample from the Illumina body map project (ArrayExpress accession number E-MTAB-513) and 3 normal breast tissue samples from the Gene Expression Omnibus dataset GSE52194. Level 3 gene expression data (raw read counts) were downloaded as provided for the TCGA samples. The normal samples within E-MTAB-513 & GSE52194 have been downloaded as raw sequence data (.fastq files) and processed using STAR aligner and htseqcount (see alignment and gene expression quantification section). Finally, to create the reference cohort, count data from all samples were merged and normalized using the Bioconductor package DESeq2^33^. Additionally, for use in predictor generation, the data were transformed into log2 scale after adding a constant +1.

### n-of-1 add-on preprocessing

OncoRep proceses a single patient sample by applying a “documentation by value” strategy ^34^. This uses preprocessing information gathered from the reference cohort generated from 1,057 breast cancer samples from TCGA. Generated thresholds can be applied to a subsequent RNA-Seq patient sample, which is a prerequisite for prospective use of transcriptomics data. Add-on preprocessing of a new patient sample was done utilizing the size factor method implemented in the DESeq2 Bioconductor package^33^. Raw read counts of a new patient sample were scaled using previously stored quantitative preprocessing information from the reference cohort, thus being the geometric mean of the counts from each gene across all samples in the reference cohort. To calculate the size factor (sequencing depth) of a new patient sample relative to the reference, the quotient of the counts in the sample divided by the counts of the reference was calculated. The median of the quotients was the scaling factor for the new patient sample. Additionally, scaled read counts were transformed to log2 scale after adding a constant +1.

### Quality control

Quality control (QC) of raw RNA-Seq reads was implemented using FastQC. Basic QC statistics are listed tabularly and linked to the full report generated by FastQC. Post alignment QC included computation of insert size distribution and collecting basic RNA-Seq metrics using functionalities provided by Picard tools.

### Alignment

RNA-Seq reads were aligned to the human genome (hg19) using STAR aligner ^35^. Alignment statistics were reported in a table within the report.

### Gene expression quantification & differential expression

Gene expression quantification was done using the htseq-count function within the Python HTSeq analysis package, which counts all reads overlapping known exons using hg19 annotation from UCSC (v57). To reduce the number of genes that serve as input for differential expression calling and pathway analysis we introduced the measure of gene expression reliability. Instead of using a non specific filtering step, a gene was determined to be reliably expressed when its expression value succeeded an expression cutoff. The expression cutoff was calculated based on the background distribution of all genes that were not expressed (raw read count equals 0) in the reference cohort (n=156 genes). This method has been described by Warren et al.^36^ and adopted for our use case. Differential expression was calculated based on a model using the negative binomial distribution as implemented in the DESeq2 package^33^.

### Prediction of receptor status & molecular subtype

Using prediction analysis for microarrays37, predictors for breast cancer receptor status (ER, PR, HER2) and molecular subtype (Luminal A, Luminal B, Her2, Basal) were implemented using samples and clinical data provided by TCGA. TCGA samples were randomly split up into a training cohort, on which the predictors were trained, and a validation cohort, on which to validate the predictors:

ER+ Training n=600; validation n=305; number of genes: 26; overall error rate training: 0.065; overall error rate validation: 0.036
PR+ Training n=600; validation n=302; number of genes: 28; overall error rate training: 0.133; overall error rate validation: 0.099

HER2+ Training n=136; number of genes: 12; overall error rate training: 0.139

Subtype Training n=346; validation n=100; number of genes: 254; overall error rate training: 0.248; overall error rate validation: 0.218

### Pathway analysis

Pathway analysis was implemented using Signaling Pathway Impact Analysis (SPIA) on the list of differentially expressed genes and their log fold changes identified in the patient sample to identify significantly dysregulated pathways using the Bioconductor packages SPIA^13^ and Graphite^38^. Graphite was used to create graph objects from pathway topologies derived from the Biocarta, KEGG, NCI and Reactome databases, which were then used with SPIA to run a topological pathway analysis.

### Fusion gene identification

Fusion gene identification was implemented using FusionCatcher^14^. FusionCatcher searches for novel/known fusion genes, translocations, and chimeras in RNA-seq data from diseased samples. The oncogenic potential of the detected fusion genes was predicted using OncoFuse^39^.

### Variant calling, filtering & annotation

Variant calling was implemented using SNPiR, a highly accurate approach to identify SNPs in RNA-seq data^15^. Basic genetic information was annotated using SnpEff^20^ and information provided by dbNSFP^21^. Variants were further filtered based on being described as either common/no known medical impact in the NCBI variants database or having a MAF *>*0.1 in the 1000 genomes data. Identified variants were further annotated using information obtained from the following databases: the Sanger Institute’s COSMIC (Catalogue of Somatic Mutations in Cancer) version 68^22^; NCBI’s ClinVar^23^; CADD (Combined Annotation Dependent Depletion) version 1.0^24^; DrugBank version 4.0^25^; and PharmGkb’s Variant and Clinical Annotations Data^26^. Entries from these databases that exactly matched the mutated allele of a single nucleotide variant, which was called by the pipeline, were included as annotations. In addition, functional effect predictions (driver or passenger status and its likely implication in the cancer phenotype) were calculated by the IntOGen^27^ pipeline and included for each variant.

### Integrative drug matching

A list of all FDA approved compounds was extracted and integrated with information from DrugBank and KEGG Drug databases, which including meta information about gene targets, pathway involvements and type of drug (e.g. inhibitor, antibody, antagonist, agonist). Altered genes were matched against these data using the meta information to select appropriate drug-gene partners. Furthermore, variants were matched against SNP-drug relationships available from DrugBank and PharmGkb.

## URLs

OncoRep: https://bitbucket.org/sulab/oncorep

Omics Pipe: https://bitbucket.org/sulab/omics_pipe

The R suite: http://www.r-project.org/

Bioconductor: http://bioconductor.org/

knitr: http://yihui.name/knitr/

knitr bootstrap: https://github.com/jimhester/knitrBootstrap

FastQC: http://www.bioinformatics.babraham.ac.uk/projects/fastqc

Picard tools: http://picard.sourceforge.net/

HTSeq: http://www-huber.embl.de/users/anders/HTSeq/doc/overview

FusionCatcher: https://code.google.com/p/fusioncatcher

OncoFuse: http://www.unav.es/genetica/oncofuse.html

SNPiR: http://lilab.stanford.edu/SNPiR

SnpEff: http://snpeff.sourceforge.net

Intogen: http://www.intogen.org ClinVar: http://www.clinvar.com DrugBank: http://www.drugbank.ca

Cosmic: http://cancer.sanger.ac.uk/cancergenome/projects/cosmic

PharmGKB: https://www.pharmgkb.org

The Cancer Genome Atlas Data Portal: http://tcga-data.nci.nih.gov/tcga

## Acknowledgements

This work was supported by the National Center for Advancing Translational Sciences (Grant UL1TR001114). The authors thank Brian Leyland-Jones, Nicholas Schork, Casey Williams, Brandon Young, Tristan Carland and Ali Torkamani for comments and assistance.

## Author contributions

T.M. designed the research, developed OncoRep and wrote the manuscript. K.F. participated in designing and developing OncoRep and wrote the manuscript. L.G. coded the variant annotation part of OncoRep. A.S. designed and supervised the research and participated in writing the manuscript.

---

**Table 1:** FDA Approved Therapies (in patients tumor type) Diff: arrow indicates if target is up- or downregulated. Mut: if checked, drug targets known mutation. Fus: if checked, drug targets fusion. PW: if checked, target is a member of altered pathway

| Target | Drugs | Diff | Mut | Fus | PW |
| --- | --- | --- | --- | --- | --- |
| DNMT1 | Decitabine | ↕ | ☐ | ☐ | ☐ |
| ERBB2 | ado-trastuzumab emtansine Pertuzumab | ↕ | ☐ | ☐ | ✓ |
| GNRHR | Degarelix | ↕ | ☐ | ☐ | ☐ |
| KCNH2 | Doxazosin | ↕ | ☐ | ☐ | ☐ |
| MMP12 | Marimastat | ↕ | ☐ | ☐ | ☐ |
| MS4A1 | Tositumomab | ↕ | ☐ | ☐ | ☐ |
| POLA1 | Fludarabine Nelarabine | ↕ | ☐ | ☐ | ✓ |
| RRM2 | Hydroxyurea | ↕ | ☐ | ☐ | ✓ |
| TOP2A | Teniposide Idarubicin | ↕ | ☐ | ☐ | ✓ |

**Table 2:**
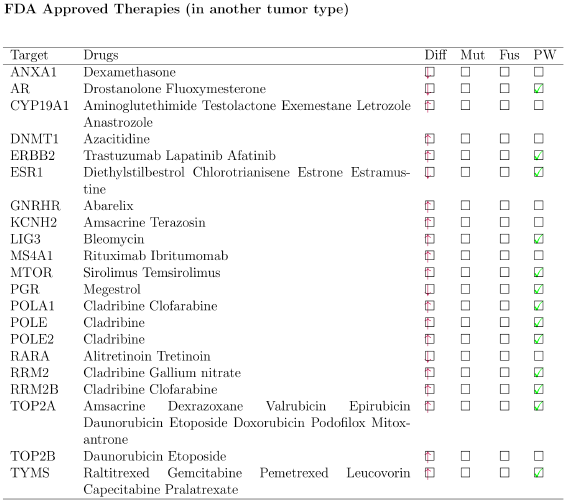
FDA Approved Therapies (in anohter tumor type) Diff: arrow indicates if target is up- or downregulated. Mut: if checked, drug targets known mutation. Fus: if checked, drug targets fusion. PW: if checked, target is a member of altered pathway

